# Nanopores reveal the stoichiometry of single oligo-adenylates produced by type III CRISPR-Cas

**DOI:** 10.1101/2023.08.18.553839

**Authors:** David Fuentenebro-Navas, Jurre A. Steens, Carlos de Lannoy, Ben Noordijk, Dick de Ridder, Raymond H.J. Staals, Sonja Schmid

## Abstract

Cyclic oligoadenylates (cOAs) are small second messenger molecules produced by the type III CRISPR-Cas system as part of the prokaryotic immune response. The role of cOAs is to allosterically activate downstream effector proteins that induce dormancy or cell death, and thus abort viral spread through the population. Interestingly, different type III systems have been reported to utilize different cOA stoichiometries (with 3 to 6 adenylate monophosphates). However, so far, their characterization has only been possible in bulk and with sophisticated equipment, while a portable assay with single-molecule resolution has been lacking. Here, we demonstrate the label-free detection of single cOA molecules using a simple protein nanopore assay. It sensitively identifies the stoichiometry of individual cOA molecules and their mixtures from synthetic and enzymatic origin. To achieve this, we trained a convolutional neural network (CNN) and validated it with a series of experiments on mono- and polydisperse cOA samples. Ultimately, we determined the stoichiometric composition of cOAs produced enzymatically by the CRISPR type III-A and III-B variants of *Thermus thermophilus*. Interestingly, both variants produce cOAs of nearly identical composition, and we discuss the biological implications of this finding. The presented nanopore-CNN workflow with single-cOA resolution can be adapted to many other signaling molecules (including eukaryotic ones), and it may be integrated into portable handheld devices with potential point-of-care applications.

---

Second messenger molecules and other small metabolites serve a wide variety of essential signaling, activation, and regulation purposes in the biological cell, such as spatial and temporal regulation of cellular responses, signal transduction between cell membrane and nucleus, and neuro-transmission ^1^. However, due to their small size, they are particularly difficult to detect, quantify, and study. In the present work, we focus on cyclic oligoadenylate molecules (cOAs) which play a crucial role in the type III CRISPR-Cas adaptive immune system of prokaryotes ^2,3^. This immune system evolved among bacteria and archaea to combat invading plasmids, bacteriophages, and other mobile genetic elements (MGEs) ^4–6^. It works by storing short DNA sequences of the encountered MGEs in an array of clustered, regularly interspaced short palindromic repeats, the CRISPR array^7^. During subsequent infections, the CRISPR array is transcribed, processed into short CRISPR RNAs (crRNAs, or guide RNAs), and incorporated into a single CRISPR-associated (Cas) protein or into multi-subunit protein complexes, with different modes of action (classified in several classes and types) ^8,7^. The crRNA-protein complexes then bind and degrade invading complementary MGEs.

The cOAs are second messengers produced by the type III CRISPR-Cas complex (Figure 1A). This ribonucleoprotein complex is endowed with three catalytic activities: sequence-specific RNAase activity by the Cas7 subunits, non-specific ssDNA cleavage by the HD domain of Cas10, and the ATP-cyclase activity of the palm domain of Cas10 responsible for generating the cOAs ^3,9^. The stoichiometry of these cOA molecules can vary between different hosts and CRISPR type III subtypes, but they typically contain three to six adenosine monophosphates (AMP) in a ring structure (cA_3_–cA_6_, Figure 1A) ^2,10,11^. The cOAs of the type III CRISPR response were found to activate particular families of downstream effector proteins containing an appropriate sensory domain, such as CARF (CRISPR-associated Rossman fold) and SAVED (second messenger oligonucleotide or dinucleotide synthetase (SMODS)- associated and fused to various effector domains). The cOA-binding domain of these proteins is often fused to other different domains with a wide variety of catalytic activities, such as (ribo-)nucleases and proteases ^8,12^. Activation of these enzymes by the cOAs results in a ‘secondary line of defense’ by type III systems, leading to the degradation of essential host biomolecules that induce either cellular dormancy or even cell death, preventing viral (or other MGE) propagation through the population ^13,14^.

**Figure 1:**
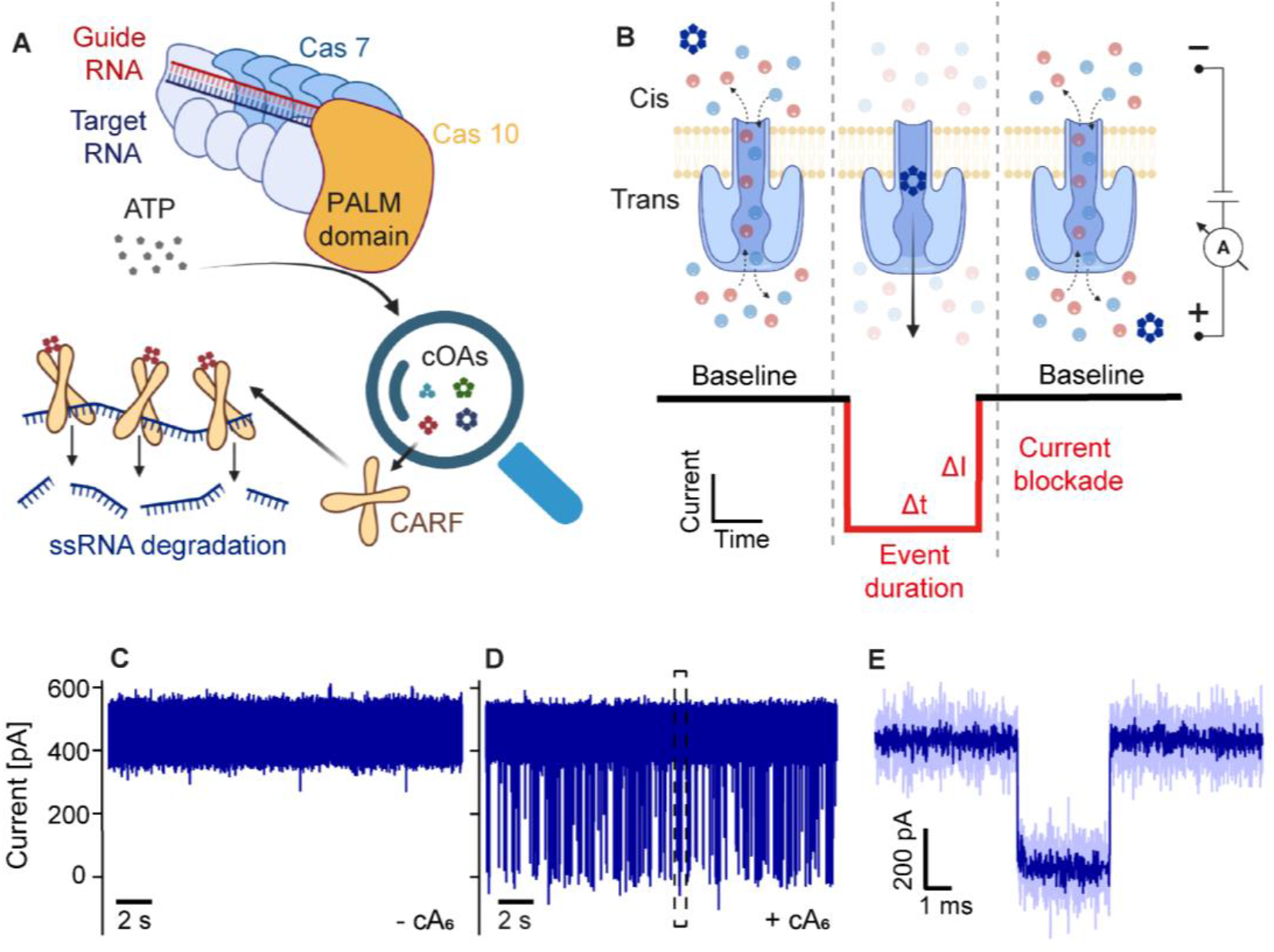
Nanopore detection of single CRISPR second messenger molecules. **A)** Schematic depiction of a type III CRISPR-Cas complex. Target RNA binding activates the cyclase activity of the palm domain of Cas10 for cOA synthesis from ATP molecules. The cOA molecules activate CARF proteins, such as the non-specific ribonuclease Csx1. **B)** Schematic arrangement of the α-hemolysin (α-HL) nanopore experiment. Upon voltage application, an ionic current flows through the pore. A cOA translocation, driven by electrophoresis, is observed as a resistive pulse with current blockade ΔI and duration Δt. We refer to the depicted pore arrangement as a ‘trans-inserted’ α-HL, where the vestibule is on the trans side. **C-E)** α-HL recordings at +200 mV in a 3 M KCl buffer, obtained as illustrated in B): **C)** Baseline trace measured without an analyte. **D)** α-HL recording after addition of 10 *μ*M cA_6_ to the cis compartment. **E)** A single cA_6_ translocation event extracted from the trace in D), as indicated by the dashed rectangle. The dark blue line represents filtered data, the light blue line represents measured raw data (see Methods).

As the diverse catalytic functions that are activated by cOAs are only just being unveiled, several questions remain unsolved. In particular: do the Cas10 homologues all produce monodisperse cOAs, or rather a polydisperse distribution? In addition, it has been reported that several distinct type III CRISPR systems (including different CARF and SAVED proteins) can be found within one host, despite using the same CRISPR array ^15–17^. For example, in *Thermus thermophilus* HB8, considered here, the genome encodes two different CRISPR-Cas type III systems, termed III-A and III-B. This is remarkable, and raises the question: what is the evolutionary benefit of having multiple distinct systems for a given task? One possible reason for the co-occurrence of multiple type III subtypes in one host is that they may each produce a unique subset of cOA stoichiometries, thus providing a regulatory benefit by activating distinct effector proteins in a fine-tuned, cOA-stoichiometry-dependent way. To test this hypothesis and elucidate cOA-dependent regulation mechanisms, a simple and rapid method to directly detect small amounts of enzymatically produced cOAs, and even quantify their stoichiometric composition with single-molecule resolution would be instrumental.

Here, we demonstrate the label-free detection of single cOA molecules and their stoichiometries using nanopore experiments, where a pore protein embedded in a free-standing lipid bilayer acts as a sensor for single cOA molecules (Figure 1B). An applied positive voltage drives the negatively charged cOA molecules through the nanopore by electrophoresis. While translocating, the cOA partially blocks the ionic through-pore current, resulting in a characteristic current blockade signal (cf. resistive pulse sensing) ^18^. Nanopores are by definition single-molecule sensors due to their small size, which we chose here to be comparable to the cOA molecule. More generally, nanopore technology is best known for commercial devices offering inexpensive and portable DNA sequencing with long-reads that are revolutionizing the life sciences ^19,20^. The sensitivity of protein nanopores is remarkable: even single enantiomers (chiral variants) in small-molecular racemates can be distinguished ^21,22^. In nanopore signal processing, neural networks play a crucial role ^23–25^. An important advantage of neural networks is their ability to implicitly extract features from the data provided during training, in contrast to other machine learning approaches that require the manual definition of informative features (e.g. hidden Markov models). Neural networks are, therefore, able to learn and combine more subtle characteristics, such as the shapes of blockade events and their current fluctuations ^26,27^. For nanopore signal processing, this enabled improved quantitative analyses ^24^. In this study, we use a convolutional neural network (CNN) to quantitatively infer the stoichiometric composition of cOA mixtures – including samples from enzymatic origin.

Our results present the first detection of cOAs with single-molecule resolution, in a sensitive nanopore assay. We compare a range of synthetic cOAs with known monodisperse stoichiometries of three to six adenosine monophosphate (AMP) monomers (subsequently: cA_3_ to cA_6_). Using this calibration data as a training set for our CNN, we turn to cOA mixtures of know polydisperse composition, and validate the capacity of the CNN to quantify the correct ratio of stoichiometries involved. We then use this new label-free cOA identification pipeline to study enzymatic cOA samples. Specifically, we identify the stoichiometric composition of cOAs produced by different type III-A and III-B complexes, and compare them with their CARF activation capability. Lastly, we discuss the implications of our results on the hypothesized evolutionary benefit of multiple type III CRISPR systems (possibly producing varied cOA stoichiometries) in one prokaryotic host. Altogether, our label-free cOA identification assay has proven its capacity to identify small second messengers with single-molecule resolution – even from enzymatic mixtures – and also their stoichiometry. This makes it a powerful tool for metabolic research of CRISPR-Cas and beyond, where fast, label-free, and quantitative readouts matter.

## RESULTS AND DISCUSSION

### Detecting single cyclic oligo-adenylates with protein nanopores

Figure 1 shows the capacity of the alpha-hemolysin (α-HL) pore protein to detect single cOA molecules. In contrast to the clean α-HL baseline measured in the absence of cOA molecules (Figure 1C), characteristic resistive pulses appear after cOA addition to the ‘cis’ side of the pore (see definition in Figure 1B, and data in Figure 1D), each representing a single cOA translocation (Figure 1E). As expected, the cOA event rate is concentration dependent (Figure S1), and the event duration decreases with increasing voltage (Figure S2, Table S1), indicating that cis-to-trans translocations take place (rather than cis-to-cis escapes). This behavior is expected for substantially charged molecules, like the poly-nucleic cOAs, for which the electrophoretic driving force dominates over other contributions (e.g. electro-osmosis) ^28^. The current blockade ΔI and event duration Δt depend on the specific analyte and measurement conditions. For cOAs with a stoichiometry of six AMP subunits (cA_6_), we find mean values and standard deviations of ΔI = 400 ± 30 pA and Δt = 315 ± 26 *μ*s, measured at +200mV in 3M KCl (Figure 1D, E, Table S1).

For the single-molecule resolution of these small cOA molecules, the choice of the nanopore and precise experimental conditions are crucial, as several factors affect the sensing performance. For example, while the MspA nanopore with its much pointier constriction site provides better spatial resolution than α-HL in DNA sequencing applications ^29,30^, it barely resolved cOA translocations (Figure S3), which limits its utility for our application in small-molecule sensing. In contrast, the α-HL with its long narrow stem yields single-molecule observations that are well resolved in time, even for these small cOA molecules. In addition, we found that the translocations through a trans-inserted α-HL (illustrated in Figure 1B) were more uniform than the translocations with a cis-inserted pore (Figure S4). Similar findings have previously been attributed to less heterogeneous excursions and interactions in the wide pore lumen of α-HL ^31^. Overall, a trans-inserted α-HL in 3M KCl and +200mV bias provided an optimal signal-to-noise ratio to resolve single cA_6_ molecules.

### Nanopore event durations reveal the stoichiometry of cOA molecules

Encouraged by the successful label-free single-molecule detection of cA_6_, we probed the sensitivity of our assay to detect even smaller cOAs: cA_3_, cA_4_, and cA_5_ depicted in Figure 2A-D. Biologically, this is highly relevant, since type III CRISPR-Cas complexes have been reported to produce cOAs with varying stoichiometries^32^. Experimentally, however, it has been impossible up to now to resolve these differences among *single* cOAs, given their small, cyclized structure (Figure S5 shows a 3D representation). Gratifyingly, with the presented nanopore assay, all cOAs down to cA_3_ can be resolved at +120mV, as evident from the current traces and zoomed-in events shown in Figure 2A-D. The measured event durations scale with the cOA size, as the larger cOAs have a reduced translocation probability through α**-**HL’s 1.4 nm constriction site (Figure 2E,F, Figures S6, S7, Table S1). Interestingly, however, cA_3_ to cA_6_ all cause similar current blockades, i.e. the larger cOAs do not measurably block more ionic current (Figure 2G,H, Figures S6, S7), likely due to local positioning inside the nanopore and 3D folding effects (cf. Figure S5). To distinguish cOA stoichiometries, we therefore focus on the distinct event durations. Their clear trend can serve as a proxy for cOA identification, while it is also evident that cA_3_ and cA_4_ cannot yet be distinguished based on the overlapping event durations alone (Figure 2E, bottom panel). We therefore turned to neural networks, which have the ability to recognize more subtle patterns in the nanopore translocation events and allow the identification of cOA stoichiometries at the single-event level.

**Figure 2:**
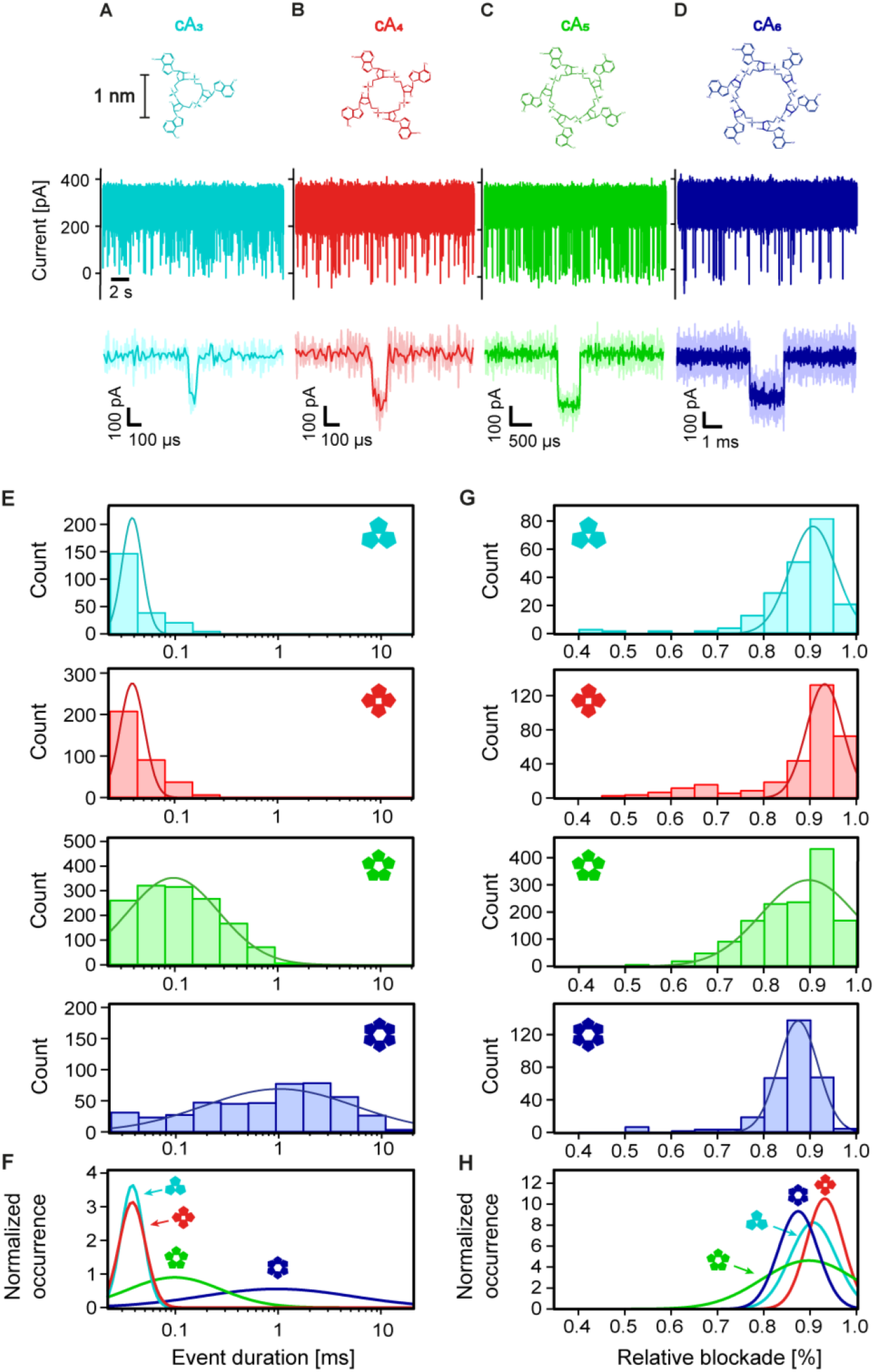
Nanopore recordings reveal the stoichiometry of cyclic oligo-adenylates. **A-D)** Nanopore recordings showing the label-free detection of single cOA molecules with different stoichiometries: cA_3_, cA_4_, cA_5_, cA_6_, respectively. Top: molecular structure of the cOAs with 3-6 AMP monomers as indicated (same scale bar for A-D). Center: nanopore current trace with short blockade events indicating cOA translocations, obtained at +120 mV with 10 *μ*M of the respective cOAs (same scale bar and axis for A-D). Bottom: zoom-in on a single translocation event of the respective cOA. **E)** Event duration histograms for the four cOAs with lognormal fits to the data. **F)** Overlay of all distributions in E), with normalized integrals. **G)** Relative blockade histograms for the four cOA datasets in E) with normal fits to the data. **H)** Overlay of the distributions in G) with normalized integrals.

### Neural networks can infer the stoichiometry of single cOA molecules

Neural networks have proven very useful in nanopore signal interpretation^33^, thanks to their ability to recognize features beyond the mere event duration and current blockade discussed above. Hence, to differentiate between nanopore events caused by cOAs of different stoichiometries, we trained a convolutional neural network (CNN) for single-event classification (Figure 3A). The training data consisted of cOA events obtained from mono-disperse cOA samples with known stoichiometry (see Methods), and the data used for model evaluation was obtained from different experiments than the training data, to achieve valid accuracy estimates. Indeed, our CNN outperforms a more conventional machine learning approach (a k-nearest neighbor classifier considering only event duration and current blockade, Figure S8), which suggests that the CNN recognizes additional signal properties, such as event shape and current fluctuations. Nevertheless, the events of the two smallest cOAs (i.e. cA_3_ and cA_4_, with very similar duration and current blockades) are not well separated by the CNN. We note that this can likely be solved, in the future, using alternative (possibly engineered) protein nanopores, capable of distinguishing both stoichiometries. For this study however, we move on with a joint class of cA_3/4_. Using this approach, the CNN correctly identified 83%, 64%, and 70% of (unseen) mono-disperse cA_3/4_, cA_5_, cA_6_ events, respectively (Figure 3B). Erroneous classifications occur mainly between adjacent stoichiometries (e.g. cA_6_ misclassified as cA_5_, or cA_5_ misclassified as cA_6_ or cA_3/4_), and mainly towards lower stoichiometries (e.g. cA_5_ misclassified as cA_3/4_, rather than cA_6_). The latter reflects the event duration distributions and their overlap (Figure 2): cA_5_ and cA_6_ events misclassified as cA_3/4_ are marked by short event durations (Figure S9). Altogether, 73% of all unseen events in monodisperse samples are correctly identified by the CNN.

**Figure 3:**
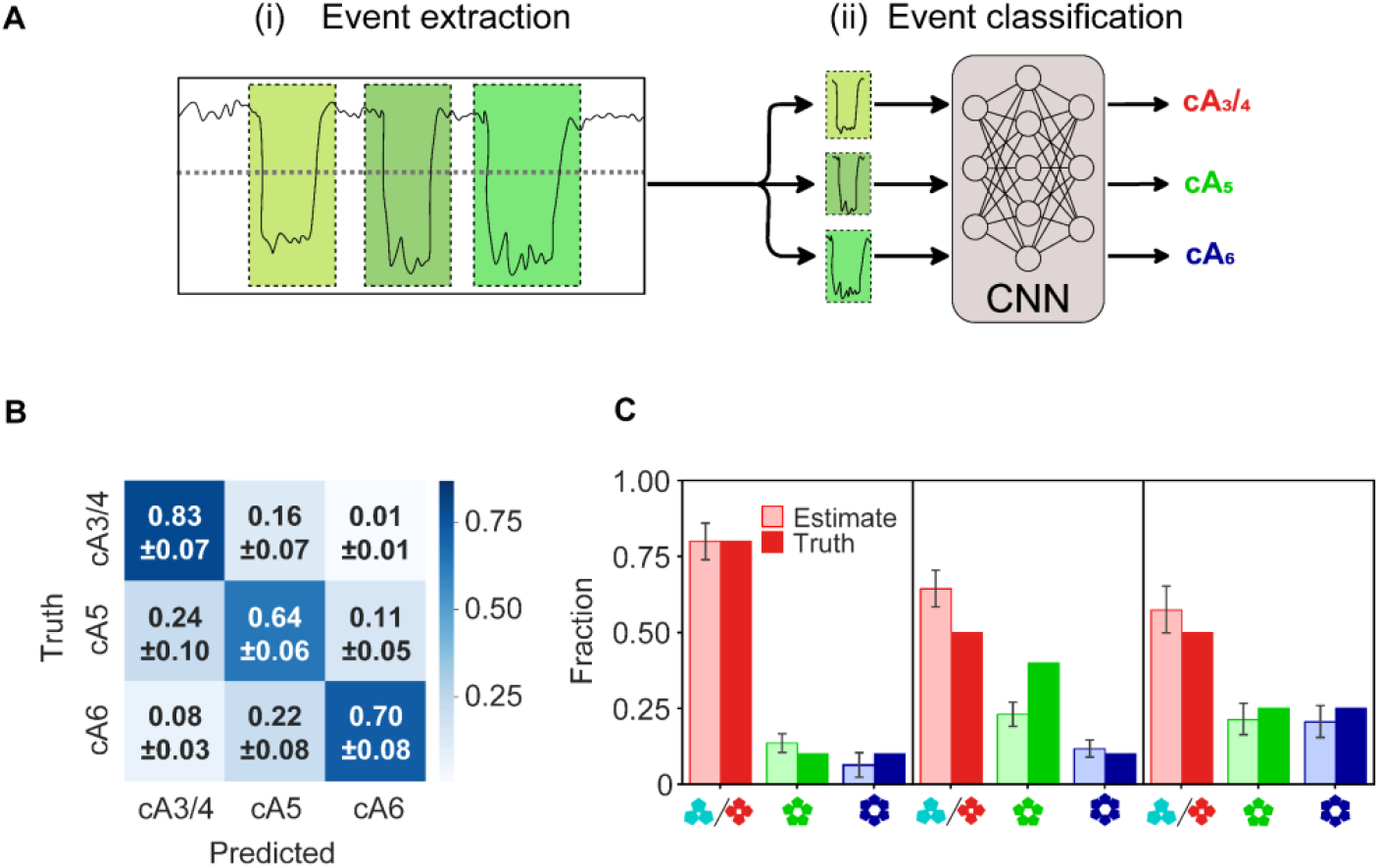
Neural network workflow and validation of the cOA stoichiometry inference from nanopore recordings. **A)** The cOA classification pipeline consists of (i) extraction of the cOA translocation events, (ii) cOA-classification per-event using a convolutional neural network (CNN). See Methods for details and code. **B)** Confusion matrix showing the performance for the single-event classification of 2,058 cOA translocation events (not included in the training set). Fractions denote relative prediction frequencies per ground truth class (mean ± standard deviation over 10 cross-validation folds). **C)** cOA stoichiometry distribution analysis: comparison of the experimental input (dark-colored) vs. the inferred output (light-colored) obtained for cOA mixtures of known composition, as indicated (percentages from left to right: 80:10:10, 50:40:10, and 50:25:25). Error bars denote 95% prediction intervals obtained by 10-fold cross validation, which express uncertainty introduced by data variability given this CNN architecture and fitting procedure.

### Quantifying polydisperse cOA mixtures and CNN validation

We next assessed the ability of the trained CNN to estimate the stoichiometric composition of polydisperse samples. We prepared three cOA mixtures with known composition, and acquired nanopore recordings of each one. Figure 3C shows the true and inferred stoichiometry fractions, with prediction intervals obtained by 10-fold cross validation (see Methods). In qualitative terms, the relative abundances (high vs. low) are correctly identified in all cases, down to the smallest tested fraction of 10%. Quantitatively, the cA_3/4_ population is partially overestimated, while the cA_5_ population is sometimes underestimated, which is consistent with our results for the mono-disperse cOAs (previous section). For the 50:40:10 mixture (center in Figure 3C), the deviation is larger than the model’s prediction interval, indicating additional imperfections beyond the training and test data variability.

Importantly, all ground truth differences in population size were correctly identified by the CNN, which inferred significantly different populations in these cases (t-test, p << 0.01, Table S2). Similarly, identical ground truth populations are also inferred to be identical – with the exception of the smallest populations of 10%. This indicates the resolution limit of the CNN classification procedure for population sizes, which we (conservatively) estimate to be 15%. Individual uncertainties for the cOA identification procedure can be estimated as 17% for cA_3,4_, 36% for cA_5_, and 30% for cA_6_, based on the confusion matrix (1 – diagonal value in Figure 3B). Equipped with this nanopore-CNN workflow with validated accuracies and uncertainty estimates, we moved on to study cOA mixtures of enzymatic origin.

### The stoichiometry of cOAs produced by type III-A and III-B CRISPR-Cas complexes

We next turned to biological cOAs, produced *in vitro* by two different CRISPR-Cas variants – type III-A and III-B – that coexist in *T. thermophilus HB8* (see Methods). If the benefit of having not one but two different type III subtypes in one organism is to activate different CARF proteins, then their cOA composition should differ. We directly tested this hypothesis with nanopore experiments using cOAs produced *in vitro* by reconstituted type III complexes ^15,34^. First, we checked if other substances present in the enzymatic reaction mix, such as ATP or short RNAs (i.e. the guide RNA and the target RNA), would interfere with the experiment. However, as they do not produce detectable nanopore signals (Figure S10), they do not interfere with the cOA quantification. Likely, the cyclized structure of the cOAs is essential for their detection, whereas the short non-cyclic RNA molecules cannot be resolved because they translocate faster through the nanopore – too fast, in fact, for the time resolution of our experiment (100kHz, cf. Figure S10). Interestingly, we find that the two CRISPR-Cas subtypes produce nearly identical cOA distributions (Figure 4A,B). The comparison with the monodisperse calibration data (Figure 4C) shows that cA_3/4_ are the predominant stoichiometries in both cases. Using the pre-trained CNN workflow, we quantified the relative abundance of cA_3/4_ as 89±16% and 81±14% for type III-A and III-B, respectively (inferred fraction ± identification uncertainty, see last section). Only minor amounts of cA_5_ (8±3% and 12±5%), and cA_6_ (3±1% and 7±3%) were detected for type III-A and III-B, respectively, which lie at the resolution limit of our technique. In absolute numbers, this converts to 870 ± 420 cOA molecules produced per type III-A complex, and 2800 ± 840 cOAs per type III-B complex under the conditions used (see Methods). In summary, the presented label-free nanopore-CNN workflow revealed that the cOA produced by the Cas10 subunit of *T. thermophilus* type III-A and III-B complexes are very similar, and they predominantly consist of cA_3/4_ (80-90%).

**Figure 4:**
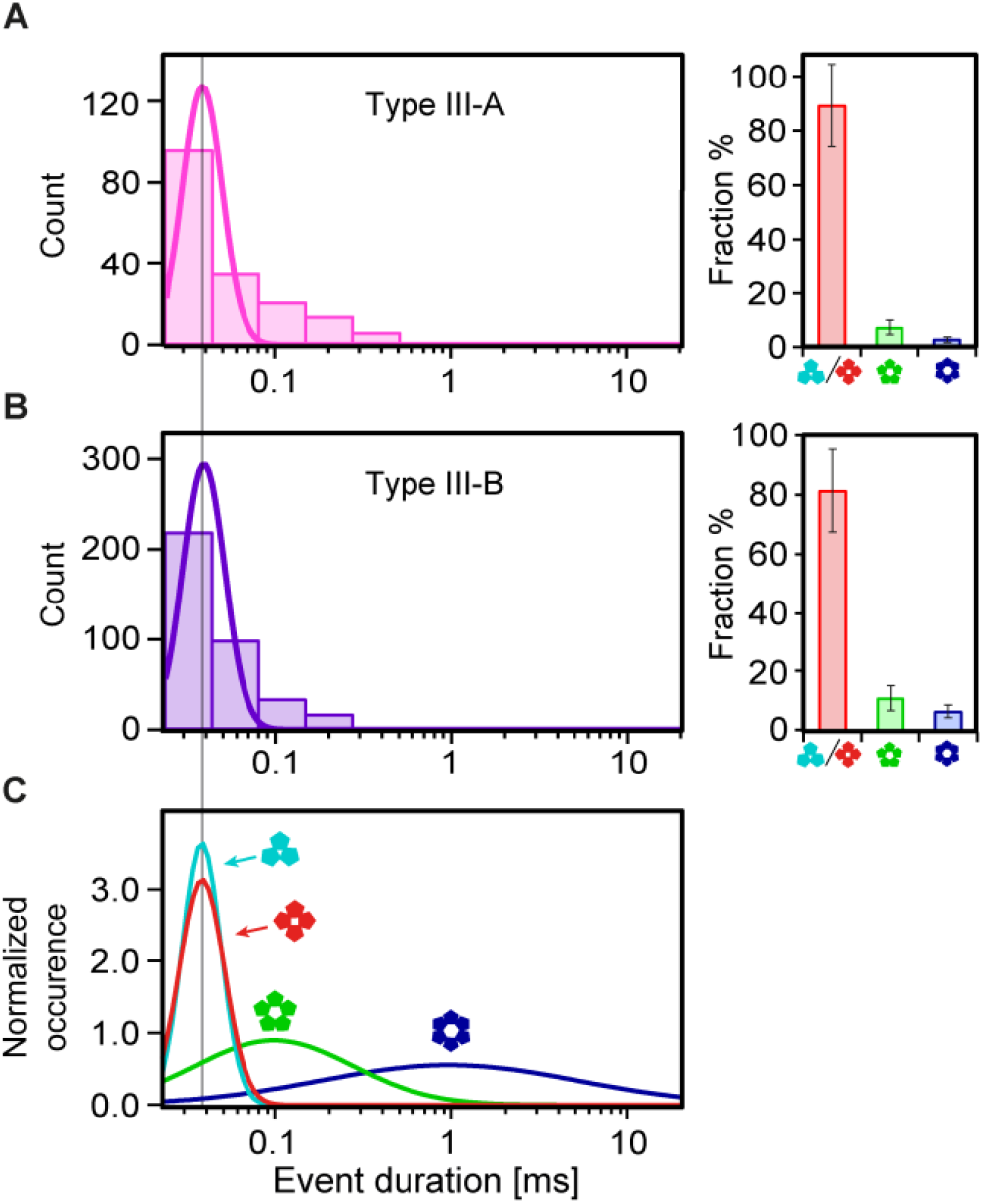
Identification of the cOA composition produced by CRISPR-Cas type III-A and III-B. **A)** Histograms of the nanopore event durations measured for the cOAs produced by the type III-A variant (measurement conditions as in Figure 2A-D). The CNN-based quantification of the stoichiometric composition is shown on the right. Error bars denote the identification uncertainty of each cOA class, estimated as 1 – the diagonal of the confusion matrix in Fig. 3B. **B)** Same as A) but for the cOAs produced by the type III-B variant. **C)** Calibration data obtained with monodisperse cOA (cf. Figure 2E). A vertical line through panels A-C provides a guide to the eye.

### The cOAs produced by CRISPR-Cas type III-A and III-B activate a cA_4_-specific CARF

After establishing that the type III-A and III-B complexes of *T. thermophilus* produce similar cOAs, we moved on to test their capacity for downstream activation of the CARF protein Csx1 from the same host (TtCsx1). Since this CARF protein has non-specific RNase activity^35^, a fluorescent RNA-cleavage reporter system was used to screen the activation of CARF proteins by cOAs from synthetic and enzymatic origin (Figure 5A, Methods). As expected^36^, among the synthetic cA_3_ to cA_6_, only cA_4_ led to TtCsx1 activation, resulting in RNA cleavage (Figure 5B). For the cOAs produced by the type III-A and III-B variants (Figure 5C), we found that both activate the RNase activity of TtCsx1 much beyond the negative controls (no target, NT). Hence, we can conclude from Figure 5C that both variants (type III-A and III-B) must indeed produce cA_4_, while other stoichiometries cannot be excluded based on Figure 5C alone. However, together with the nanopore-CNN results above, our results show that both type III variants produce a nearly identical cOA composition, including cA_4_. These results argue against the hypothesis that individual Cas10 homologues produce distinct cOA compositions for a homologue-specific downstream regulation in *T. Thermophilus*. The presented nanopore-CNN workflow has thus elucidated the hitherto unknown stoichiometric composition of the cOA second messengers of these type III-A and III-B systems, and given the results, the purpose of multiple CRISPR-Cas homologues in one host remains an open question.

**Figure 5:**
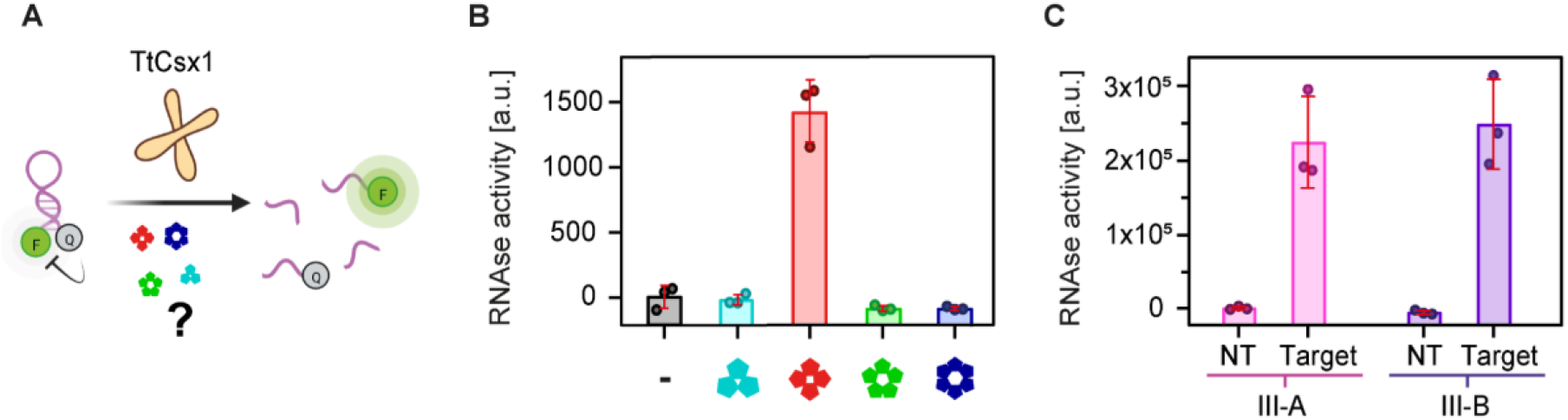
The CARF activation by enzymatic and synthetic cOAs reveals stoichiometric specificity. **A)** Schematic representation of the RNA cleavage assay. Reporter RNA molecules conjugated with a fluorophore (F) and a quencher (Q) are incubated with the CARF RNase TtCsx1 and various cOA samples. If the cOAs activate TtCsx1, it cleaves the reporter RNA, leading to increased fluorescence intensities compared to the negative controls. **B)** Activity assay of TtCsx1 after the addition of synthetic monodisperse cOAs, or no cOAs in case of the negative control (-). Assays were run in triplicates. Also, cA2 was tested and did not activate TtCsx1 (Figure S11). **C)** TtCsx1 RNase activity upon addition of cOAs produced in an *in vitro* reaction mixture by endogenous type III-A and III-B from *T. thermophilus*, as indicated. cOAs are only produced in the presence of complementary target RNA (Target). Reactions with non-complementary target RNA (no target, NT) serve as a negative control for the assay, run in triplicates.

## CONCLUSION

We present a cOA identification workflow with single-molecule resolution (Figure 6) that combines the potential of nanopore technology, neural networks, and enzymatic assays. For the first time, we individually detected these small cOA second-messenger molecules from the type III CRISPR-Cas systems using nanopores. Based on the observed nanopore event durations, we were able to distinguish cA_3/4_ from cA_5_, and cA_6_. Using monodisperse synthetic cOAs, we acquired calibration data for each stoichiometry, and used it to train a CNN, which we validated using cOA samples of known polydisperse composition. We then detected cOAs of enzymatic origin, produced by the type III-A and type III-B CRISPR-Cas complexes of *T. thermophilus*, and quantified the previously unknown stoichiometric composition using the validated CNN. Additional enzyme activity assays revealed the stoichiometry-specific activation of CARF proteins, and further complement the nanopore-CNN results.

**Figure 6:**
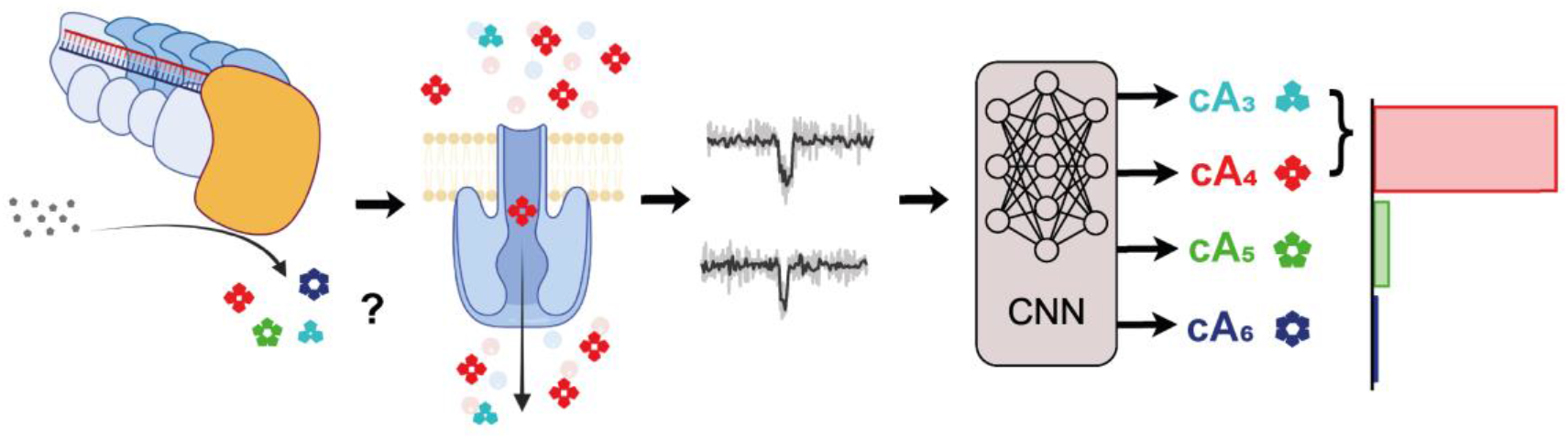
The nanopore-CNN pipeline for the identification of single second messengers from type III CRISPR-Cas. Biological cOAs of unknown stoichiometry, produced by CRISPR type III-A and III-B complexes, are detected one-by-one in nanopore experiments. Single-molecule events extracted from the nanopore reads are then identified by a pre-trained CNN, yielding the stoichiometric composition of the biological cOA samples.

We conclude from the nanopore experiments that (i) cyclic oligo-adenylates (cA_3_ to cA_6_) can be sensitively detected at the single-molecule level; (ii) the cA_3/4_, cA_5_, cA_6_ stoichiometries cause distinguishable nanopore event durations; and (iii) the type III-A and type III-B CRISPR-Cas complexes produce cOAs of qualitatively near-identical stoichiometry: predominantly the small cA_3_ or cA_4_. Using a pre-trained CNN, we (iv) identify the stoichiometries of the cOA at the single-event level with an accuracy of 73%, and we (v) quantify the cOA composition produced by *T. thermophilus* CRISPR-Cas type III-A and type III-B variants as 89±16% and 81±14% cA_3/4_, respectively, and only trace amounts of cA_5_ and cA_6_ in both cases. We also estimate (vi) the absolute number of cOAs produced which amounts to 870 ± 420 cOAs per type III-A complex, and 2800 ± 840 cOAs per type III-B complex, under the conditions used. Lastly, (vii) the enzymatic RNA cleavage assays verified that the combined cA_3,4_ class must contain cA_4_, proving that CRISPR-Cas type III-A and III-B both produce this specific second messenger. Altogether, we thus find nearly identical stoichiometric cOA compositions in both variants, which challenges the hypothesis that co-occurring type III complexes would produce distinct cOA for specific downstream regulation purposes.

In the future, the presented nanopore-CNN pipeline can be used to elucidate the cOA stoichiometries of different CRISPR-Cas type III, and it is readily adaptable to additional signaling molecules, which include – but are not limited to – signaling molecules of other antiviral immune systems, such as CBASS (cyclic oligonucleotide-based antiphage signaling system)^37^. Additional protein nanopores will be explored to detect also cA_2_ and to distinguish cA_3_ from cA_4_, which was not possible with our current hemolysin nanopore. Promising candidates include (possibly engineered) pore proteins with a long narrow channel (hemolysin, aerolysin ^38^), or with multiple constrictions (CsgG ^39,40^). Altogether, we demonstrated the first single-molecule sensor for CRISPR second messengers. We achieved stoichiometry sensitivity using affordable nanopore technology that can be integrated into handheld devices and may, in the future, enable quantitative point-of-care diagnostics^41^ with single-molecule resolution.

## Methods

### Protein nanopore recordings

were performed using the Montal-Mueller technique ^42^, using an Axopatch 200B amplifier and a Digidata 1550B digitizer (both Molecular Devices) with a sampling rate of 500kHz, and low-pass filtered at 100kHz. A free-standing lipid bilayer was formed in a custom-built flowcell with two buffer reservoirs separated by a Teflon film (GoodFellow, Huntingdon, England) with a small electro-sparked aperture (ca. 100*μ*m diameter, obtained with a spark generator, Daedalion, Colorado, United States). The Teflon film was pre-treated on each side with 10*μ*l of a hexadecane solution (10% hexadecane, Acros Organics, Geel, Belgium) in pentane (Alfa Aesar, Massachusetts, United States), and let dry for 5 minutes. Both reservoirs were filled with 400*μ*l of measurement buffer each (3M KCl, 100mM Tris, 0.1 mM EDTA, pH 8), connected to Ag/AgCl electrodes (silver wire with 0.5mm diameter, Advent, Oxford, England, chloridized in household bleach), and the flowcell was placed inside a Faraday cage. Bilayers were built by adding 10*μ*l of a 1,2-diphytanoyl-sn-glycero-3-phosphocholine solution (DPhPC, 10mg/ml, Avanti Polar Lipids, Alabama, United States) in pentane onto each buffer reservoir and pipetting up and down as described previously ^42^. α-HL oligomers (kindly provided by Sergey Kalachikov, Columbia University) were added to the trans reservoir (with the working electrode under positive polarity) and their spontaneous membrane insertion caused a characteristic current. The cOA solution (500*μ*M, Biolog, Bremen, Germany) was added to the cis reservoir (with the ground electrode) to a final concentration of 10*μ*M.

### Nanopore data processing

was performed in Igor Pro (v6.37, Wavemetrics, Oregon, United States) using custom code. Event detection was performed after filtering the signal with a digital 80 kHz low-pass filter and median-conserving decimation to a final sampling rate of 80kHz (25*μ*s time resolution). Events were extracted by applying a threshold at 65% of the open-pore current. The mean current blockades were calculated from the extracted data. To calculate event durations, 1,000-fold bootstrapping with replacement was performed, where each subset was fit with a single-exponential (with X offset). The uncertainty is expressed as the standard deviation of all bootstrapped time constants.

### Neural networks: per-event classification

To classify single cOA translocation events, we trained a 1D convolutional neural network (CNN) implemented using Tensorflow ^43^. As training data, we used 48 traces of +120 mV translocations from synthetic monodisperse cOA samples with known stoichiometries. Events were extracted as described above, individually normalized and scaled by the standard deviation. To provide information about the relative blockade, we include 47 datapoints (5,875 ms) of the baseline signal on either side of the extracted event. Moreover, to ensure consistent input size, we pad each to a width of 250 datapoints with zeroes on the right-hand side. The original event duration is concatenated to this signal as a final feature.

We feed each individual padded event into the CNN, which performs multi-class classification. The CNN consists of two 1D convolutional layers with 10 filters of width 25 and a ReLu activation function, each of which is followed by a batch normalization and 20% dropout operations. Next, values are maxpooled with a pool size of 10, and fed into a dense layer with 4 output nodes (one per cOA class) and a softmax activation function. The CNN is trained for 100 epochs at a learning rate of 0.001 with the Adam optimizer that minimizes cross entropy loss based on the 3 one-hot encoded cOA classes. During training, we perform oversampling to prevent class imbalance. Five restarts were performed, after which the classifier with the highest training accuracy was selected for testing. Training was done on a laptop equipped with an Intel Core i7 processor (8 cores @3GHz, Santa Clara, CA) and a T500 GPU (NVIDIA, Santa Clara, CA). Training runtime was around 20 minutes.

### CNN validation

was done using a 10-fold cross-validation scheme. Importantly, events of a given trace were never split between training and test sets, thus ensuring that no trace-specific characteristics were learned. The reported overall accuracy was calculated over the predictions of all folds merged into one data set. We compared the CNN performance to a traditional feature-extraction-based classifier by performing k-nearest neighbor (KNN) classification based on relative blockade and (log transformed) dwell times (k=3 after hyperparameter optimization), implemented using sci-kit learn ^44^. In cOA mixture classifications, *t*-tests and two one-sided *t*-tests (TOST) procedures were used to test difference and equivalence respectively between all cOA distributions. The TOST procedure tests whether two cOA relative distributions *N*(*μ*_*cAx*_, *σ*_*cAx*_)and *N*(*μ*_*cAy*_, *σ*_*cAy*_) have means diverging less than a given level *δ*, by performing two one-sided t-tests, with null-hypotheses:

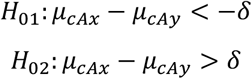

Where rejection of both *H*_01_ and *H*_02_ means that the alternative hypothesis that the means differ less than *δ* must be accepted, or:

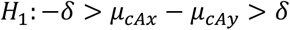

Both methods are implemented in python using the scipy package ^45^ and available in the main code repository for this paper (see Code availability).

### The abundance estimation

ability was evaluated by running inference using the trained model on polydisperse samples of known composition. To obtain prediction intervals reflecting variation induced by training data, prediction was repeated with all 10 classifiers obtained during cross-validation. We compensate for differences in event rate by using cOA-specific constants which convert from event count to estimated abundance in a sample. The constants are estimated by counting the number of events in the training files, and normalizing for concentration and time. The values of these correction factors (in units of events s^-1^ *μ*Mol^-1^) were 0.17±0.05, 0.62 ±0.7 and 0.12 ±0.3 for cA_3/4_, cA_5_, and cA_6_ respectively.

### Enzymatic cOA synthesis and quantification

Either TtCsm (type III-A) or TtCmr (type III-B) endogenous protein complexes from *Thermus thermophilus* (purified as described previously ^15,34^) were incubated in 150 mM NaCl, 20 mM Tris-HCl pH 8, 10 mM DTT, 2 mM MgCl and 1 mM ATP in a total volume of 20 *μ*l, to which 200 nM of non-target or target RNA (IDT, Coralville, Iowa) was added, as indicated. The reactions were carried out in triplicate by incubating the samples for 60 minutes at 65°C. Target RNA sequence:

5′ GAACUGCGCCUUGACGUGGUCGUCCCCGGGCGCCUUAUCUACGGCCAUCG 3′.

Non-target RNA sequence:

5′ UGAUGAGGUAGUAGGUUGUAUAGUAAGCUUGGCACUGGCCGUCGUUUACG 3′.

The absolute numbers of each cOA produced by the type III-A and III-B complexes were calculated from the observed nanopore event rate *r*_*e*_ (in units of s^-1^), the CNN deduced fraction *cA*_*x*_% of each stoichiometry class ‘x’, and the pre-defined correction factor *CF*_*x*_ (see last section), the dilution factor *d* (between the enzyme reaction and the nanopore experiment), and the Avogadro constant *N*_*A*_:

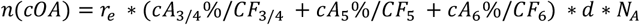

For the type III-A and III-B measurements, the respective event rates were 0.47±0.19 s^-1^ and 1.17±0.11 s^-1^, and the dilution factors were *d*=21 and *d*=27.6, respectively. We report the cOA numbers normalized per type III-A or III-B complex (their concentration during cOA synthesis was 62.5 nM), and the uncertainty was calculated by error propagation of the reported individual uncertainty estimates.

### CARF cOA stringency assay

The CARF protein Csx1 from *Thermus thermophilus*, 1*μ*M *T*tCsx1, purified as described previously ^41^, was incubated with 250 nM RNaseAlertTM (ThermoFisher, Waltham, USA) in 150 mM NaCl, 20 mM Tris-HCl pH 8, 10 mM DTT, and 1 mM ATP in a total volume of 20 *μ*l, to which 1 *μ*M of synthetic cOAn (n=2-6, Biolog, Bremen, Germany) was added. The reactions were carried out in triplicate for 60 minutes at 65ºC in a Bio-Rad CFX384TM Real-Time System (Hercules USA), measuring FAM at 1 min intervals. Data shows the relative fluorescence at 30 min.

### CARF activation assay with enzymatic cOA from type III CRISPR

Either type III-A or type III-B endogenous complexes from *Thermus Thermophilus* were incubated with 1 *μ*M TtCsx1 (the CARF protein), 250 nM RNaseAlertTM (ThermoFisher, Waltham, USA) in 150 mM NaCl, 20 mM Tris-HCl pH 8, 10 mM DTT, 2 mM MgCl and 1 mM ATP in a total volume of 20 *μ*l, to which 200 nM of non-target or target RNA (IDT, Coralville, Iowa) was added, as indicated. The reactions were carried out in triplicate for 60 minutes at 65ºC in a ThermoFisher QuantStudio 1 Real-Time PCR System (Waltham USA). Data shows the relative fluorescence at 30 min.

Target RNA sequence: GAACUGCGCCUUGACGUGGUCGUCCCCGGGCGCCUUAUCUACGGCCAUCG.

Non-target RNA sequence: UGAUGAGGUAGUAGGUUGUAUAGUAAGCUUGGCACUGGCCGUCGUUUACG.

## Supporting information

SI_Fuentenebro

## Code availability

CNN training and evaluation code is freely available on Gitlab at https://github.com/cvdelannoy/coa_classifier

## Notes

J.A.S. is a founder and shareholder of Scope Biosciences B.V. R.H.J.S is a shareholder and members of the scientific board of Scope Biosciences B.V. J.A.S and R.H.J.S are inventors on CRISPR-Cas related patents/patent applications.

## Acknowledgments

We thank Sergey Kalachikov and Jens Gundlach for kindly providing aHL and MspA, respectively. We thank Chenyu Wen for critical reading of the manuscript, and Giovanni Maglia for early discussions on instrumentation. R.H.J.S. is supported by a VIDI grant (VI.Vidi.203.074) from The Netherlands Organization for Scientific Research (NWO). SS acknowledges IPM4 funding from Wageningen University, and the NWO XL grant ProPore: OCENW.XL21.XL21.003.

