## Supplementary material for "Nanopores reveal the stoichiometry of single oligo-adenylates produced by type III CRISPR-Cas": SI_Fuentenebro

by Fuentenebro-Navas et al.

### Table of contents:

**Figure S1:** The concentration-dependent cOA event rates (2).

**Figure S2:** The voltage-dependent event rates and durations (2).

**Figure S3:** The MspA nanopore detects cOAs with shorter event durations than α-HL (3)

**Figure S4:** Comparison of the cis vs trans-inserted α-HL nanopore regarding its cOA detection capacity (4).

**Figure S5:** cOAs have a variable dimeter in solution (5).

**Figure S6:** Aligned nanopore translocation events of cOA (6).

**Figure S7:** 2D scatter plot of the nanopore translocation events of the indicated cOA molecules (7).

**Figure S8:** k-nearest neighbor performance on cOA event classification (8).

**Figure S9:** Event duration distributions of unseen nanopore events, per ground truth and CNN-predicted stoichiometry (9)

**Figure S10:** Control data confirming that neither ATP nor RNA cause detectable nanopore events, while cOAs are detected in Type IIIA/B samples (10).

**Figure S11:** cA2 does not activate TtCsx1

**Table S1:** Nanopore event durations of cA3-6 at different voltages (11).

**Table S2:** Pair-wise statistical difference and equivalence test p-values, for neural network classifications of cOA mixtures (12).

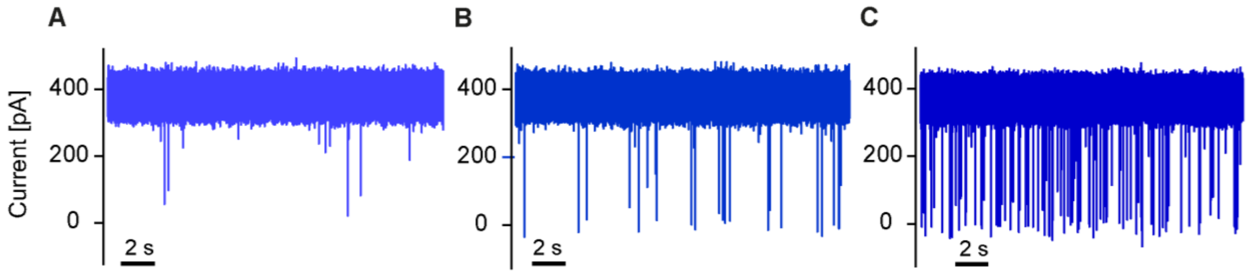

**Figure S1: Concentration-dependent cOA event rates.** Current traces recorded with a trans-inserted α-HL pore at +160mV in a 3M KCl buffer (Methods) after the addition of $cA_6$ to the cis chamber: **A)** 100nM, **B)** 1µM, **C)** 10µM $cA_6$. The event rates are: 0.2, 0.7, and 3.8 events/s, respectively. $cA_6$ events are also detectable below 100nM, albeit at correspondingly slower event rates.

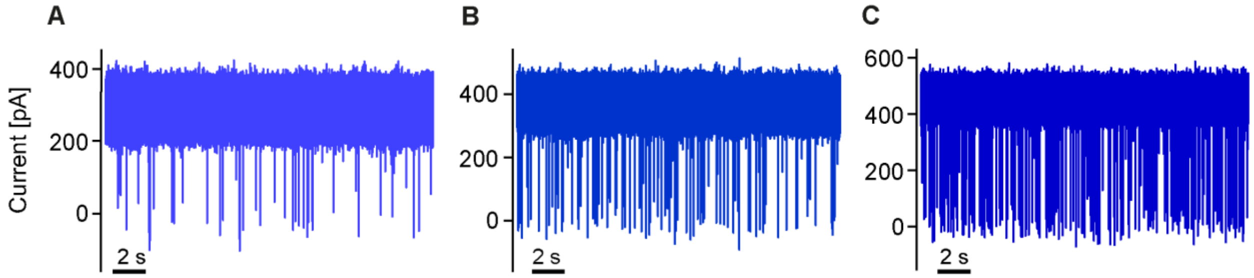

**Figure S2: Voltage-dependent event rates and durations.** Current traces recorded with a trans-inserted α-HL pore in a 3M KCl buffer (Methods) after the addition of 10µM $cA_6$ to the cis chamber, recorded at **A)** +120mV, **B)** +160mV, **C)** +200mV. As expected for electrophoresis-driven translocation events, the event rate increases with voltage (1.7, 4.3, and 7 events/s, respectively) while the event duration decreases. The voltage-dependent event durations for $cA_3$, $cA_4$, $cA_5$, $cA_6$ are specified in **Supplementary table 1**.

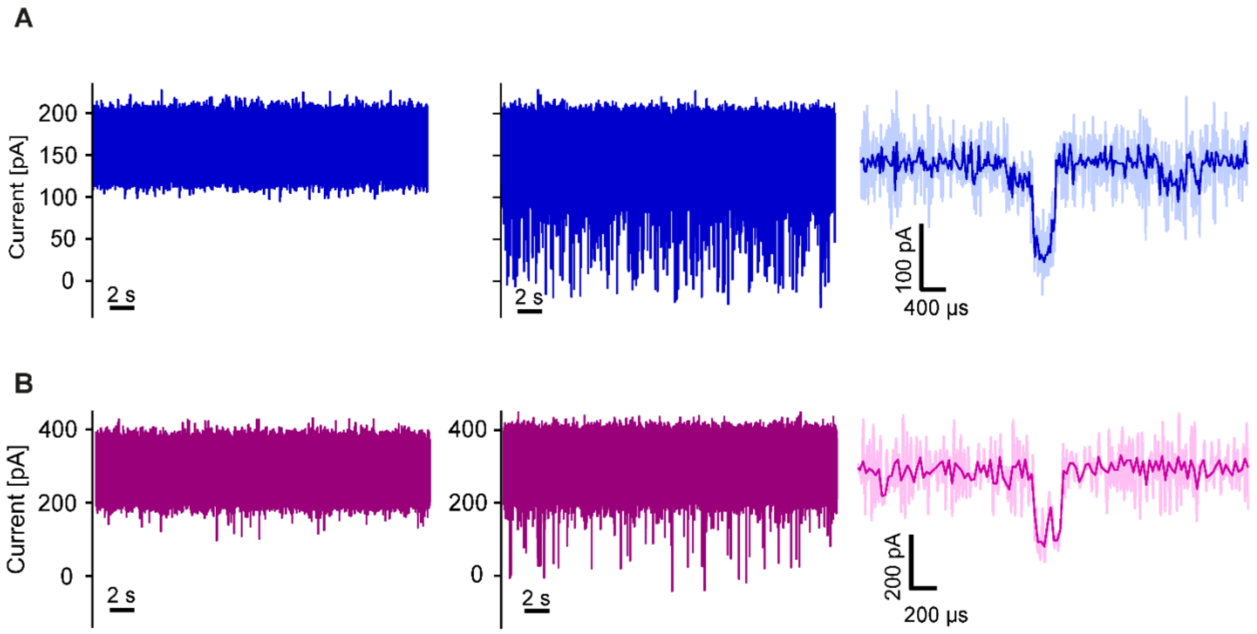

**Figure S3: The MspA nanopore detects cOAs with shorter event durations than α-HL**. **A)** α-HL. **B)** MspA. Both A, B) were measured under identical conditions: a cis-inserted pore at +160mV in a 1M KCl buffer (Methods). Left: baseline without analyte. Middle: current trace recorded after the addition of 10µM $cA_6$ to the cis chamber. Right: zoom-in on a single event. The events detected with MspA are often truncated and generally shorter than those with α-HL, making the latter a more suitable choice for cOA detection.

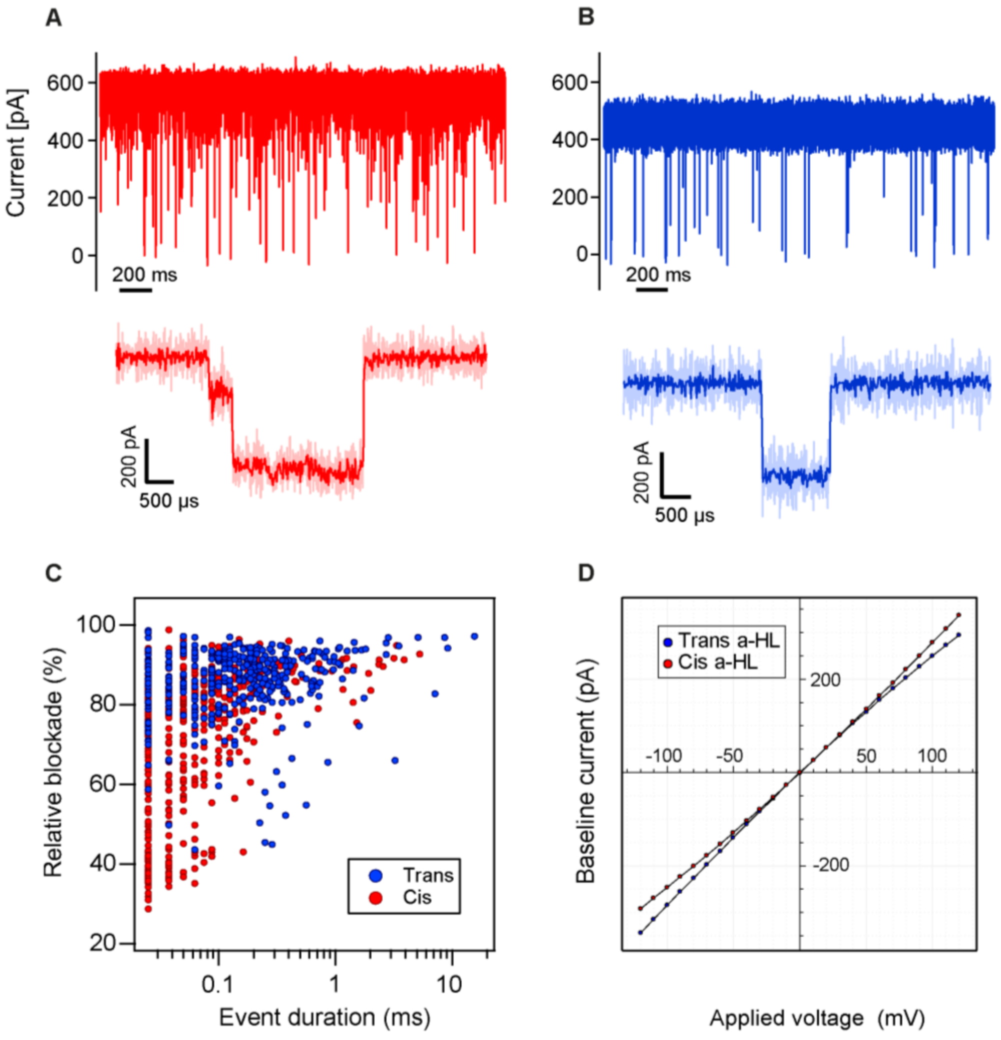

**Figure S4: Comparison of the cOA detection capacity of cis vs. trans-inserted α-HL nanopores. A,B)** show a current trace and event zoom-in recorded with a cis- (A) or trans-inserted α-HL pore (B), respectively, both measured at +200mV, in 3M KCl, with 10µM $cA_6$ in cis. The events observed with a cis-inserted α-HL were less uniform. **C)** Scatter plot of blockade vs. duration for the events cis in A and trans events in B, respectively. The events of the cis-inserted α-HL (red) are more heterogenous (spread out) than the more uniform ones of the trans-inserted pore (blue). **D)** Current-voltage curves of cis- and a trans-inserted α-HL pores as indicated (without cOA).

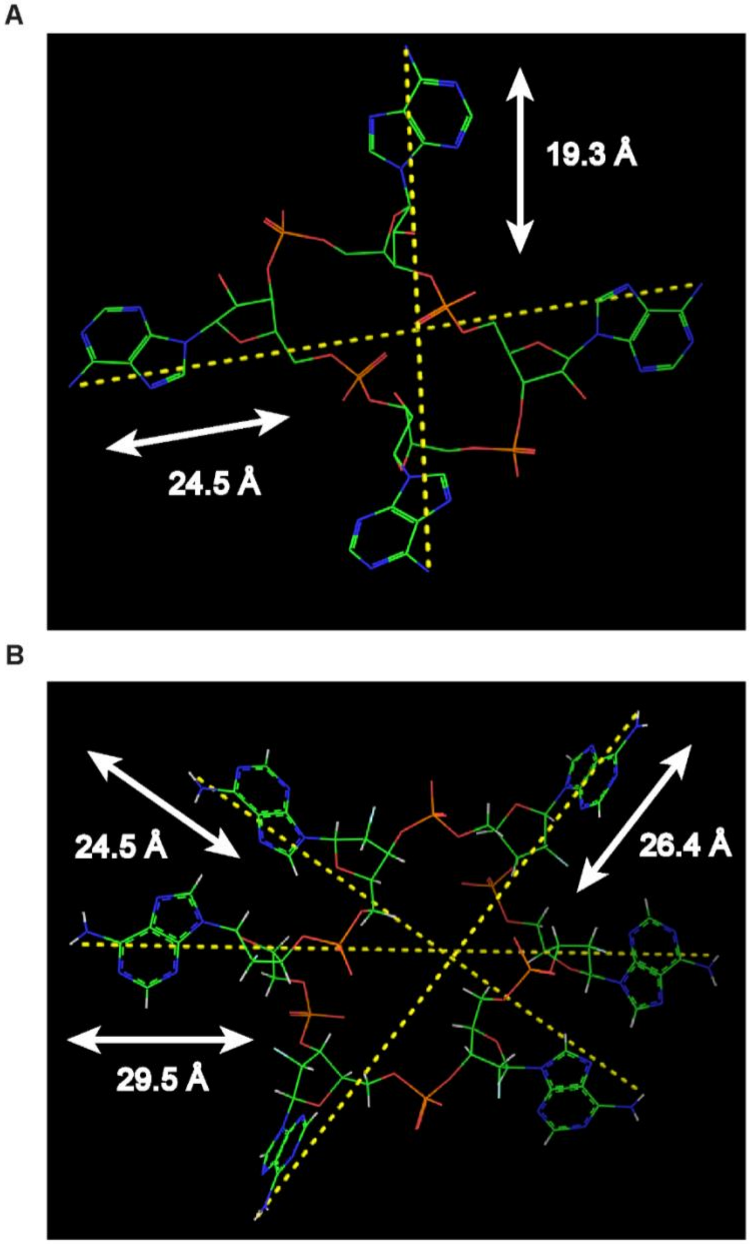

**Figure S5: cOAs have a variable diameter in solution.** 3D structures of $cA_4$ and $cA_6$ in complex with **A)** SsCsa3 (PDB ID: 6W11) and **B)** EiCsm6 (PDB ID: 6TUG) proteins, respectively. The proteins have been hidden for visualization purposes. The cOA have flexible bonds and fold in solution. This leads to a variable diameter, which allows them to squeeze through the narrower α-HL pore (1.2 nm). Note that the molecule shown at the bottom is cyclic hexa-2'-fluoro-hexa-dAMP ($cFA_6$), a $cA_6$ mimic resistant to degradation but still able to activate its substrate EiCsm6.

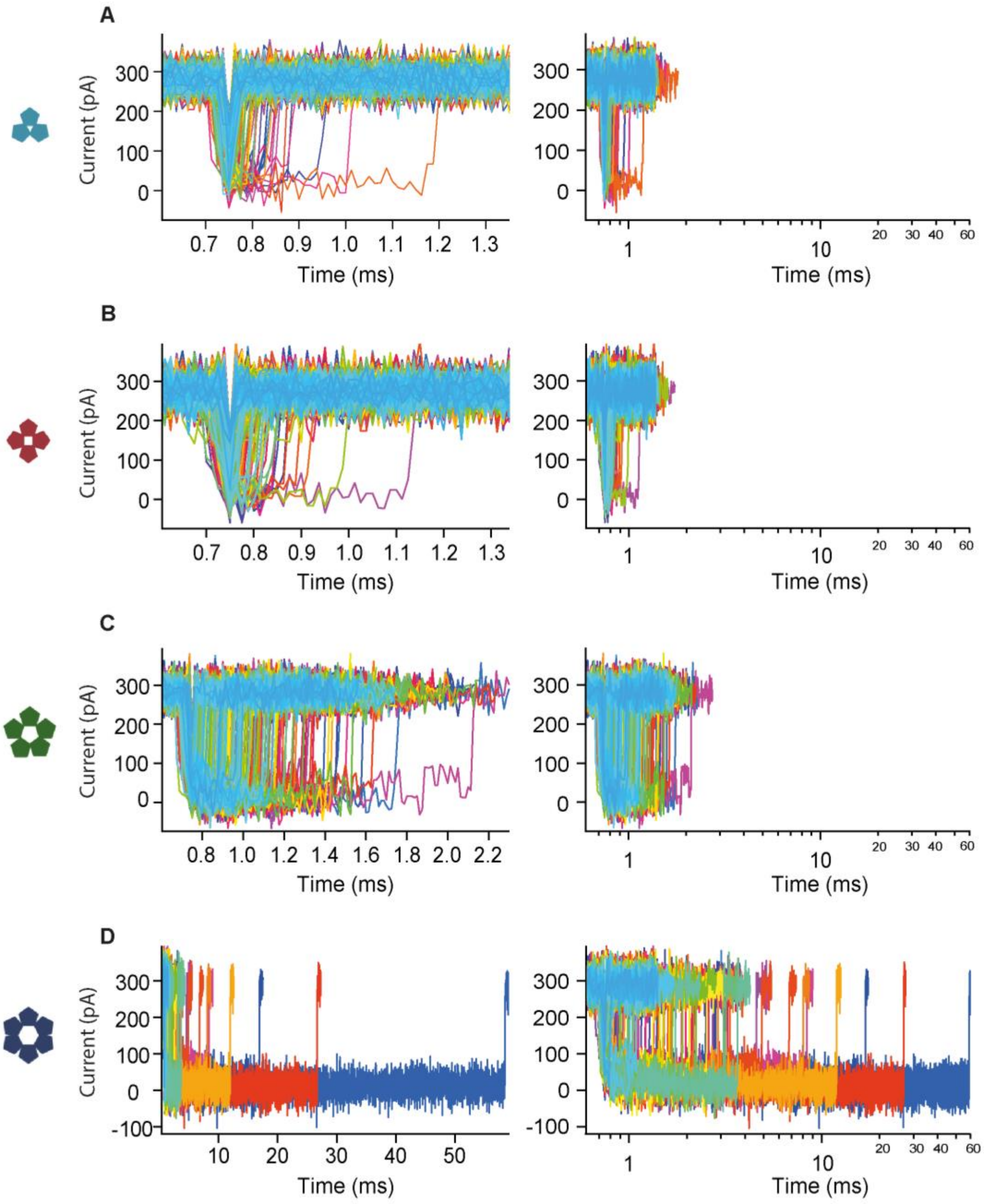

**Figure S6: Aligned nanopore translocation events of cOAs: A) $cA_3$, B) $cA_4$, C) $cA_5$, D) $cA_6$.** The events were obtained as in Figure 2A (10µM cOA in cis chamber, trans-inserted α-HL pore, +120mV, 3M KCl buffer). The events are aligned at their starting point and individually colored. Plots with linear (right) and logarithmic (left) time axes are shown.

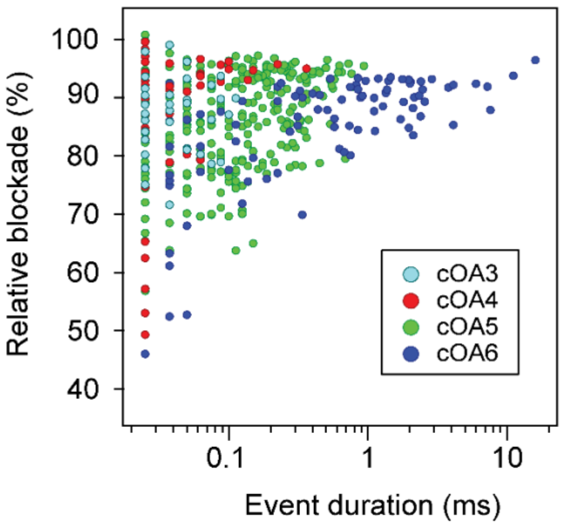

**Figure S7: 2D scatter plot of the nanopore translocation events of the indicated cOA molecules**. Each dot represents a single translocation event, represented by its relative current blockade vs. its duration.

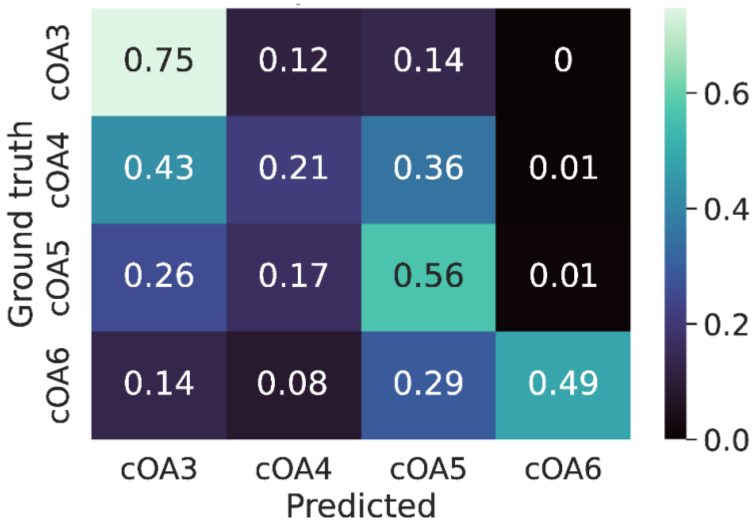

**Figure S8: k-nearest neighbor performance on cOA event classification.** This confusion matrix shows the performance of a k-nearest neighbor (kNN) classifier for cOA event identification, fed with the same data as the CNNs (Fig 3B). The kNN only uses current blockade and event duration as features for classification, and performs poorly when compared to the trained CNN. This is likely due to the fact that the CNN is able to recognize more subtle patterns present in the signal, such as event shape and current fluctuations, leading to a better classification of unseen events.

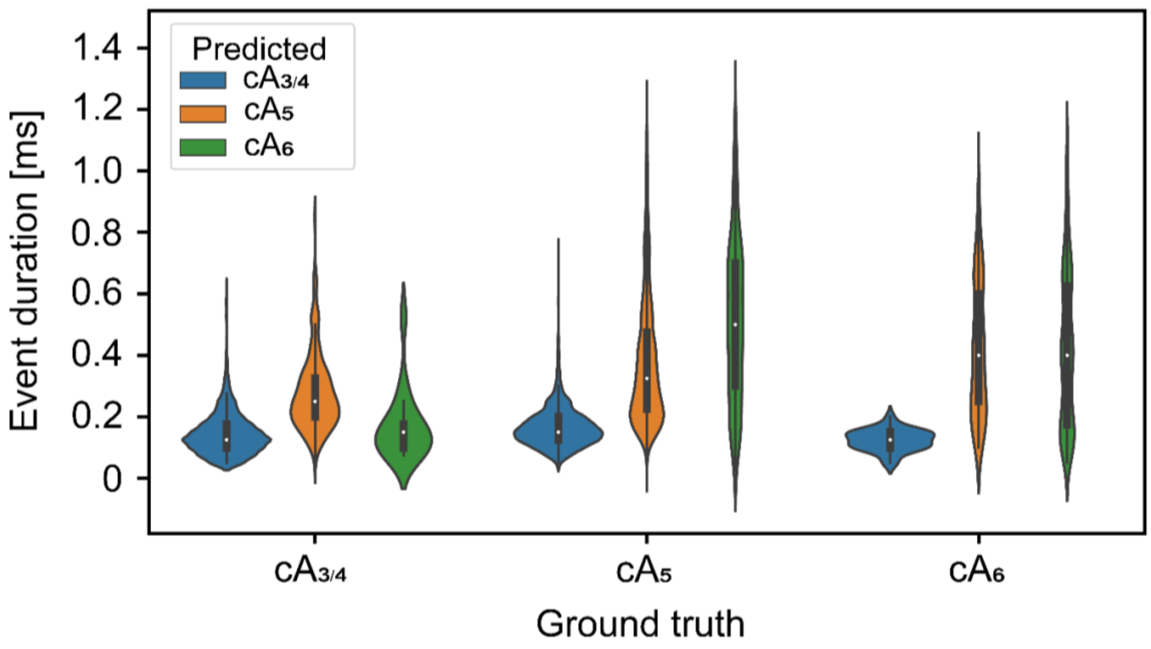

**Figure S9: Event duration distributions of unseen nanopore events, per ground truth and CNN-predicted stoichiometry.** Test events from mono-disperse samples were classified using the trained CNN and subsequently analyzed on event duration for each combination of ground truth and CNN-predicted stoichiometry. Distributions are shown as violin plots, where each violin surface is normalized to the same surface area to allow for visual distribution comparison. Events that were erroneously predicted to be of lower-stoichiometry than indicated by the ground truth showed shorter event durations, indicating that this feature played a major role in misclassifications.

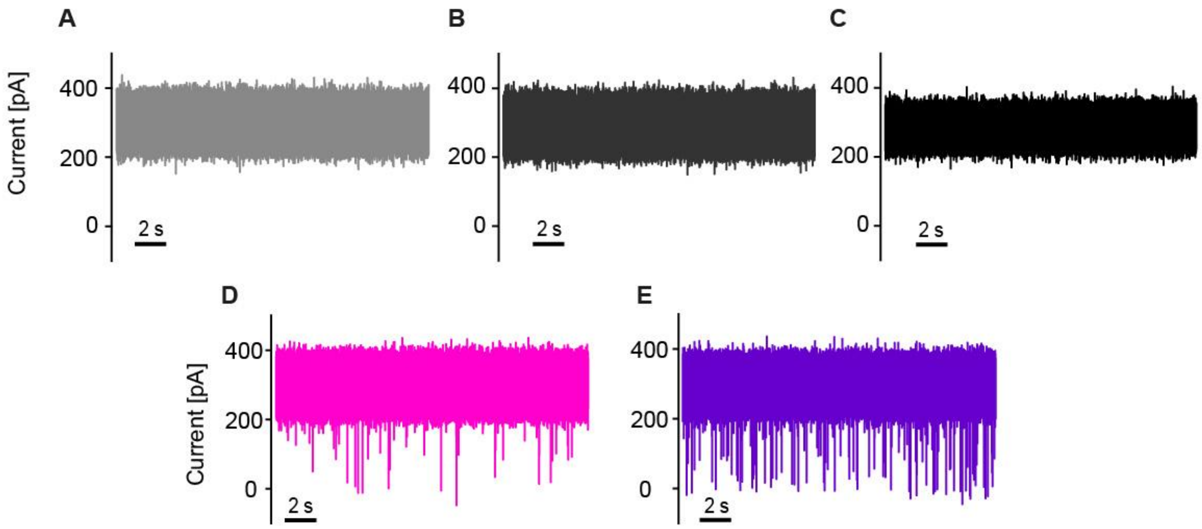

**Figure S10: Negative control: neither ATP nor RNA from enzymatic reactions cause detectable nanopore events**. All current traces were recorded with a trans-inserted α-HL pore at +120mV in 3M KCl buffer. **A)** Baseline without analyte. **B)** 1µM ATP in the cis chamber (as used in Fig. 4B,C, cf. Methods) does not cause any detectable events. **C)** The enzyme reaction mix (incl. enzyme, RNA, as used in Fig. 4B,C, see Methods) does not cause any detectable events in the *absence* of ATP (precluding enzymatic cOA synthesis). B) and C) confirm that the events detected in Figure 4B,C originate from bona-fide Cas10-produced cOAs, and not from other molecules present in the reaction. **D,E)** Current traces recorded with the *active* reaction mix (incl. ATP) show typical cOA events, indicating the presence of enzymatically produced cOA, synthesized by CRISPR/Cas type III-A (panel D) and III-B proteins (panel E), respectively.

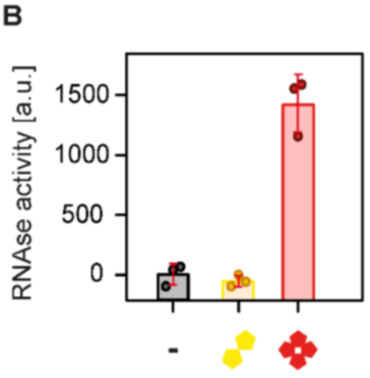

**Figure S11: $cA_2$ does not activate TtCsx1.** The RNA cleavage assay (cf. Fig. 5) was extended to $cA_2$. In contrast to $cA_4$, it does not activate the RNAse activity TtCsx1 beyond the negative control (-).

**Table S1: Nanopore event durations of $cA_{3-6}$ at different voltages.** The time constants and standard deviations (SD) were obtained by 1,000-fold bootstrapping with replacement, fitting the bootstrapped event duration histograms to an exponential function. The average tau and SD values from the 1,000 bootstrapped fits are reported. The nanopore recordings were performed as described in Figure S2, and the values were calculated from at least n=150 events. All cOAs show the same trend: decreasing event durations with increasing voltages, indicating translocations instead of trapping events. For the smaller $cA_3$ and $cA_4$, with shorter event durations, higher voltages push the event durations below the time resolution (25µs) and the events are not resolved. Altogether, a voltage of +120mV was found to provide the optimal balance between distinct event durations, capture rate, and signal-to-noise for the detection of all four cOAs tested in this work.

| | Event duration & SD (µs) | | | |
|---|---|---|---|---|
| | cA3 | cA4 | cA5 | cA6 |
| +80mV | 32±5 | 109±22 | 297 ± 33 | 2010 ± 944 |
| +120mV | 15±1 | 22± 2 | 96 ±4 | 1213 ± 119 |
| +160mV | Not resolved | 14±3 | 61 ±6 | 396 ± 39 |
| +200mV | Not resolved | Not resolved | 55 ±5 | 315 ± 26 |

**Table S2: Pair-wise statistical difference and equivalence test p-values, for neural network classifications of cOA mixtures.** t-test p-values and two one-sided t-test (TOST) procedure p-values are reported for difference and equivalence testing respectively. Nomenclature: sample '50_40_10' indicates 50% $cA_{3/4}$, 40% $cA_5$ and 10% $cA_6$. While the $cA_5$ and $cA_6$ in the 80_10_10 mixture differ significantly according to the t-test ($p \ll 0.01$), in pairwise equivalence tests, they deviate significantly less than 10% (TOST procedure, $p<0.05$). The $cA_5$ and $cA_6$ fractions differ less than 10% from each other for both samples (TOST procedure, $p \ll 0.01$).

| *Sample:* | $cA_A$ | $cA_B$ | $p_{TOST}$ | $p_{t\text{-}test}$ |
|---|---|---|---|---|
| *50_25_25* | $cA_{3/4}$ | $cA_5$ | 1 | $3.77 \times 10^{-13}$ |
| *50_25_25* | $cA_{3/4}$ | $cA_6$ | 1 | $2.71 \times 10^{-13}$ |
| *50_25_25* | $cA_5$ | $cA_6$ | $1.21 \times 10^{-5}$ | $4.88 \times 10^{-1}$ |
| *50_40_10* | $cA_{3/4}$ | $cA_5$ | 1 | $1.95 \times 10^{-16}$ |
| *50_40_10* | $cA_{3/4}$ | $cA_6$ | 1 | $2.61 \times 10^{-19}$ |
| *50_40_10* | $cA_5$ | $cA_6$ | $8.18 \times 10^{-1}$ | $1.25 \times 10^{-8}$ |
| *80_10_10* | $cA_{3/4}$ | $cA_5$ | 1 | $3.76 \times 10^{-20}$ |
| *80_10_10* | $cA_{3/4}$ | $cA_6$ | 1 | $8.54 \times 10^{-21}$ |
| *80_10_10* | $cA_5$ | $cA_6$ | $3.76 \times 10^{-2}$ | $1.33 \times 10^{-6}$ |
| *type III-A* | $cA_{3/4}$ | $cA_5$ | 1 | $1.29 \times 10^{-305}$ |
| *type III-A* | $cA_{3/4}$ | $cA_6$ | 1 | $1.44 \times 10^{-300}$ |
| *type III-A* | $cA_5$ | $cA_6$ | $3.33 \times 10^{-91}$ | $2.07 \times 10^{-74}$ |
| *type III-B* | $cA_{3/4}$ | $cA_5$ | 1 | $6.03 \times 10^{-269}$ |
| *type III-B* | $cA_{3/4}$ | $cA_6$ | 1 | $4.25 \times 10^{-261}$ |
| *type III-B* | $cA_5$ | $cA_6$ | $3.56 \times 10^{-66}$ | $2.65 \times 10^{-50}$ |
